# Distinct co-modulation rules of synaptic and voltage-gated currents coordinates interactions of multiple neuromodulators

**DOI:** 10.1101/265694

**Authors:** Xinping Li, Dirk Bucher, Farzan Nadim

## Abstract

Different neuromodulators usually activate distinct receptors but can have overlapping targets. Consequently, circuit output depends on neuromodulator interactions at shared targets, a poorly understood process. We explored quantitative rules of co-modulation of two principal targets: voltage-gated and synaptic ionic currents. In the stomatogastric ganglion of the crab *Cancer borealis*, the neuropeptides proctolin and CCAP modulate synapses of the pyloric circuit, and activate a voltage-gated current (*I*_MI_) in multiple neurons. We examined the validity of a simple dose-dependent quantitative rule that co-modulation by proctolin and CCAP is predicted by the linear sum of the individual effects of each modulator, up to saturation. We found that this rule is valid for co-modulation of synapses, but not for the activation of *I*_MI_, where co-modulation was sublinear. Given the evolutionary conservation of neuromodulator receptors and signaling pathways, such distinct rules for co-modulation of different targets are likely to be common across neuronal circuits.

## Introduction

All nervous systems adapt to changes in the environment and the internal state of the animal. In different contexts, awake or asleep, fed or hungry, light or dark, neuronal circuits produce different output (Xia and Mills, 2004; Inagaki et al., 2014; Wester and McBain, 2014; Burke et al., 2015; Filosa et al., 2016). This context-dependent output is actively shaped by various neuromodulators through changes in neuronal and synaptic properties (reviewed in Brezina, 2010; Bargmann, 2012; Marder, 2012; Nadim and Bucher, 2014). The large number of neuromodulators identified within species clearly indicates that, at any time, every neuronal circuit is co-modulated by a number of these substances (Marder and Bucher, 2007; Taghert and Nitabach, 2012; van den Pol, 2012; Richter et al., 2014; Su et al., 2014). The combination and distribution of neuromodulators present depends on context, and often is the means to convey it (Cohn et al., 2015; Lovett-Barron et al., 2017; White et al., 2017). Consequently, essential behaviors such as breathing, sleeping, learning, and mating, as well as cognitive tasks, rely on combined actions of multiple neuromodulators (Doi and Ramirez, 2008; Woods et al., 2014; He et al., 2015; Yamazoe-Umemoto et al., 2015; Mena et al., 2016; Asahina, 2017; Donlea et al., 2017). Thus, proper neuronal circuit function depends on specific combinations of neuromodulators, and how they act in concert.

While much is known about the actions of single neuromodulators, few studies have explored how multiple neuromodulators interact. Most of these studies have provided qualitative descriptions of altered output at the systems level (Brezina et al., 1996; Dickinson et al., 1997; Mesce et al., 2001; Thirumalai and Marder, 2002; Beliez et al., 2014). Only a handful of studies have explored the combined actions of neuromodulators on their direct targets, also mostly qualitatively (McCormick and Pape, 1990; Parker, 2000; Djokaj et al., 2001; Svensson et al., 2001; Park and Spruston, 2012; Garcia et al., 2015).

Neuromodulator targets fall into two categories: ionic currents that shape neuronal excitability, and synapses, which determine circuit organization. In a single neuron, a single neuromodulator can have multiple subcellular targets (divergence) and multiple neuromodulators can have overlapping targets (convergence) (reviewed in Nadim and Bucher, 2014). Such patterns of divergence and convergence can result in complex co-modulatory effects on neuron and synapse function, and consequently circuit output.

To understand how co-modulation shapes circuit output, it is important to characterize how co-modulation occurs at shared targets. Here we focus on convergent co-modulation of synapses and voltage-gated currents by exploring 1) if the combined actions of neuromodulators on a shared target can be predicted quantitatively from their individual actions, and 2) if co-modulation of synaptic and voltage-gated ionic currents in a neuron follows the same rule. For neuromodulators with converging signaling pathways, the most parsimonious prediction would be that their effects at a shared target simply add up linearly to produce a combined effect, up to the saturation level. It should be noted, however, that such linear addition does not exclude the possibility that each separate modulator effect might have a distinct dose-dependence that is inherently nonlinear. In addition, the dynamics and physiological effects of modulating a target can be complex and nonlinear.

In this study, we used the pyloric circuit of the crab stomatogastric ganglion (STG) to examine whether the dose-dependent actions of two peptide neuromodulators on their targets can be predicted by the linear summation of their individual actions, up to saturation. Several peptides activate *I*_MI_, a voltage-gated ionic current (Golowasch and Marder, 1992; Swensen and Marder, 2000) in STG neurons, likely through converging signaling pathways from different receptors (Garcia et al., 2015; Gray et al., 2017). Some also modulate pyloric synapses (Thirumalai et al., 2006; Zhao et al., 2011; Garcia et al., 2015). We measured the influence of two peptide neuromodulators on synaptic currents and on *I*_MI_. Because the influence of the peptides on these components can be assayed simultaneously, they provide a good test for understanding the rules of co-modulation of different aspects of neuronal processing. We found that co-modulation of synaptic transmission and the voltage-gated current follows distinct rules—a mechanism likely to be generalizable. The machinery underlying neuromodulation is evolutionarily well conserved and most receptors have homologs across invertebrate and vertebrate systems (Mirabeau and Joly, 2013; Lovett-Barron et al., 2017), and many neuromodulators share G-protein mediated signaling pathways (Doi and Ramirez, 2008). Thus, such distinct rules for co-modulation of different components are likely to be used in other neuronal circuits and by other neuromodulators.

## Results

We explored the modulatory effects of the two neuropeptides CCAP and Proc on *I*_MI_ in the lateral pyloric (LP), and on the reciprocal synapses between LP and the pyloric dilator (PD) and neurons. The influence of these peptides on pyloric neurons and synapses can be assayed simultaneously, while all other neuromodulatory inputs are removed.

We began by quantifying the individual modulatory effects of CCAP and Proc on both synapses and *I*_MI_ in the LP neuron across a range of concentrations, ranging from subthreshold to saturation. These dose-dependent quantifications allowed us to build predictors of the modulatory effect of each individual modulator at any concentration.

We then characterized the effect of co-application of both peptides in two stages. First, we examined if co-modulation is history dependent by co-applying the peptides following exposure to either Proc or CCAP, as interactions between neuromodulators can depend the order of application and produce priming or gating (Dickinson et al., 1997; Svensson et al., 2001). Then, in separate experiments, we tested the effect of various combinations of the two peptides, applied at different concentrations, and compared the results with the predictions of the linear summation rule.

### Dose-dependent effect of individual peptides on the synapses

We quantified the individual modulatory effects of CCAP and Proc in separate sets of experiments. In each experiment, we measured the effect of the peptide on both the LP to PD and the PD to LP synapses. Hence, we will discuss four different synapse-peptide cases: LP to PD-CCAP, LP to PD-Proc, PD to LP-CCAP and PD to LP-Proc.

In each synapse-peptide case, we measured the postsynaptic current in control and in increasing concentrations of the peptide with simultaneous two-electrode voltage clamp recordings of both neurons (Figure 1A). In the STG, two identical PD neurons and the anterior burster (AB) neuron are strongly electrically coupled and form the pacemaker group. Unless specified otherwise, the PD to LP synapse in this study refers to the combined synaptic current from the pacemaker group to the LP neuron. As expected for a graded synapse, the amplitude of postsynaptic current increased as the presynaptic step voltage increased (Figure 1B and C). The current-voltage relationship of each synapse was fit with the sigmoidal curve given by equation (1), which is described by three parameters: *I*_max_ (synaptic amplitude), *V*_mid_ (halfactivation voltage) and *V*_c_ (slope factor at *V*_mid_). A more positive *V*_mid_ indicates a higher threshold for activation and larger *V*_c_ means a shallower activation curve (Figure 1D). For each synapse-peptide pair, we examined how *I*_max_, *V*_mid_ and *V*_c_ were changed by the peptides (Figure 2).

**Figure 1.**
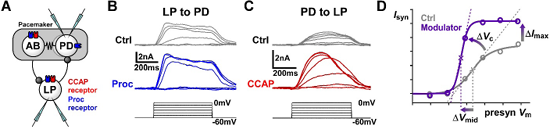
CCAP and Proc modulate the strength and activation curves of the reciprocal synapses between the LP and PD neurons. **(A)** Schematic diagram of the synaptic connectivity between the electrically-coupled (resistor symbol) pyloric pacemaker neurons, AB and PD, and the follower LP neuron. Both synapses (stick-and-ball symbols) are inhibitory. Also shown are the known receptor expression for CCAP and putative receptor expression for Proc in these neurons. The experimental protocol involved simultaneous two-electrode voltage-clamp recordings of the PD and LP neurons. **(B)** Example recordings of postsynaptic currents measured in the PD neuron in response to voltage steps in the presynaptic LP neuron in control saline (Ctrl) and in the presence of 1 μM Proc. Measurements were done in 0.1 μM TTX. **(C)** Example recordings of synaptic currents measured in the LP neuron in response to voltage steps in the presynaptic PD neuron in control saline (Ctrl) and in the presence of 1 μM CCAP. Measurements were done in 0.1 μM TTX. **(D)** To measure the modulatory effects, the mean value of the postsynaptic currents was plotted against the presynaptic voltage and fit with a Boltzmann type sigmoidal function. Changes in maximum synaptic current (*I*_max_), half-activation voltage (*V*_mid_) and slope factor (*V*_c_) were compared in control and in the presence of the modulator.

**Figure 2.**
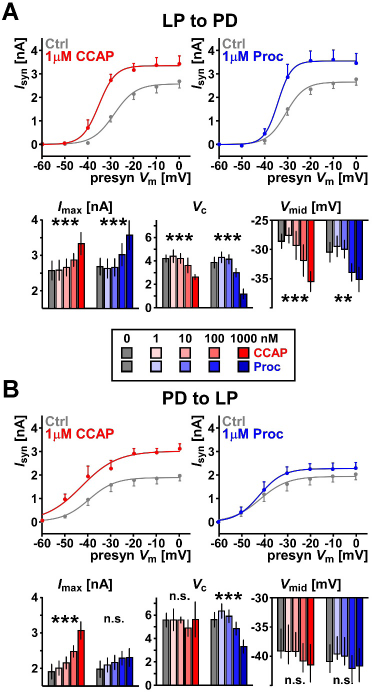
CCAP and Proc modulate the synapses between the LP and PD neurons in a dose-dependent manner. **(A)** Both CCAP and Proc increase the amplitude of the LP to PD postsynaptic current (*I*_syn_). Top panels show mean and SEM of *I*_syn_ as well as sigmoidal fits for control and modulators applied at the maximum concentration of 1 μM. As the applied concentration is increased, CCAP increases *I*_max_ (p<0.0001), decreases the slope factor *V*_c_ (p<0.001) and decreases *V*_mid_ (p<0.0001). (All tests One-Way RM-ANOVA, N=5.) Proc has a s*I*_MI_lar effect on these three parameters (p<0.0001 for *I*_max_ and *V*_c_, p=0.0047 for *V*_mid_, OneWay RM-ANOVA, N=6). **(B)** As the applied concentration increases, CCAP, but not Proc, increases the amplitude of the PD to LP synapse. Top panels as in A. CCAP increases *I*_max_ (p<0.0001), but not V_mid_ (p=0.50) or V_c_ (p=0.95), Proc modulates *V*_c_ (p<0.0001) but not *I*_max_ (p=0.22) or *V*_mid_ (p=0.11). All tests OneWay RM-ANOVA, N=6. (** p<0.01, *** p<0.001). All raw data are provided in Figure 2-source data.

For the LP to PD synapse, both CCAP and Proc significantly increased *I*_max_, shifted *V*_mid_ to more negative potentials, and reduced *V*_c_ across concentrations (Figure 2A). In contrast, for the PD to LP synapse, CCAP only increased *I*_max_, but did not affect *V*_mid_ or *V*_c_, while Proc only decreased *V*_c_, but did not affect *I*_max_ or *V*_mid_ (Figure 2A).

Notably, the same peptide differentially modulated different synapses. For example, CCAP changed *I*_max_, V_mid_, and V_c_ at the LP to PD synapse, but only *I*_max_ at the PD to LP synapse. In addition, different peptides had different effects on the same synapse. For example, CCAP changed only *I*_max_ at the PD to LP synapse, while Proc changed *V*_c_. Overall, both CCAP and Proc strengthened both synapses, although the manner of modulation depended on the synapse and the modulator.

We used the data shown in Figure 2 to build predictors for each synapse-peptide pair. The predictor is a surface fit to all synaptic current amplitudes, measured at different presynaptic voltage steps and modulation concentrations (Figure 3), which has a sigmoidal relationship with both the presynaptic voltage and the log of the modulator concentration (fit given by equation (3)). These predictors allow us to estimate the synaptic current at any voltage and modulator concentration by interpolation. The surface fits also allow us to visualize and measure the distinct modulation effects of the two peptides on each synapse and of each peptide on the two synapses.

**Figure 3.**
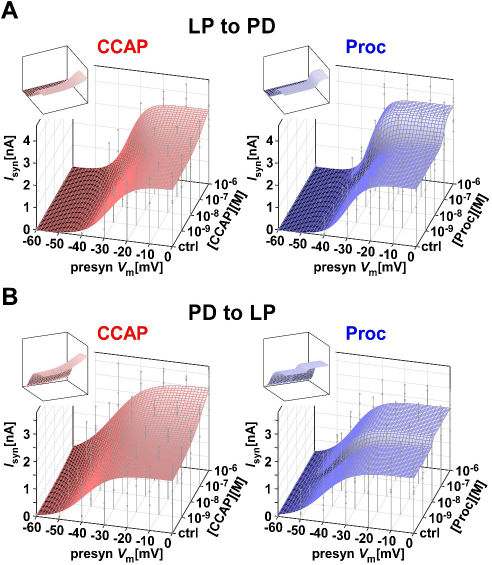
The dose-dependent influence of CCAP and Proc on the activation curves of the two synapses was used to construct predictors of modulation on synapses. **(A)** A double-sigmoidal surface fit (equation (3)) to the activation data of the LP to PD synapse in different doses of Proc or CCAP can be used to estimate the influence of the respective modulator on the synapse at any presynaptic voltage and any concentration of the modulator. Droplines indicate measurement points of the experimental data, with the filled circles marking the data points. Insets show the same surface from a different viewpoint. **(B)** Same as (A), but for the PD to LP synapse. The fit parameters were: panel A, CCAP: a1=3.619, a2=−1.042, a3=−38.00, a4=9.890, a5=3.197, a6=1.920, *C*_mid_=−6.556, *C*_c_=0.5555; panel A, Proc: 01=3.508, θ2=−0.902, θ3=−34.68, 04=4.320, 05=2.9 13, 06=1.324, *C*_mid_=−7.018, C«=0.1359; panel B, CCAP: 01=3.632, 02=−1.735, 03=−44.74, 04=5.82, 05=8.135, 06=−2.116, *C*_mid_=−6.455, *C*_c_=0.8039; panel B, Proc: 01=2.273, 02=−0.2560, 03=−42.43, 04=2.090, 05=5.184, 06=1.126, *C*_mid_=−7.958, *C*_c_=0.04605.

### The saturation level of the co-modulatory effect on the synapses is not history dependent

Our main hypothesis assumes that the saturation of synaptic co-modulation is not affected by the order of application; that is, one modulator does not gate or prime the effect of the other modulator. Prior to testing our hypothesis, it was therefore important to verify this assumption. To test if the co-modulatory saturation level depended on the prior application of either modulator, we did two separate sets of experiments for each synapse. In each experiment, we saturated the synapse with either Proc or CCAP first, and then with both peptides co-applied.

Saturation of neuromodulatory effects can occur when the receptors, the signaling pathways, or the targets themselves reach maximum capacity. Co-modulatory effects at high concentrations depend on the degree to which the different neuromodulators occlude each other's effects. If the separate effects of two neuromodulators saturate because the common target is maximally modulated, the effect of each modulator occludes the effect of the other. If the separate effects of two neuromodulators saturate because their respective receptors are saturated, neither modulator's effect should completely occlude the other's. If the signaling pathways saturate, occlusion depends on pathway interactions.

We first examined if co-modulation produced an additional effect above that of the single neuromodulator at 1 μM, the presumed saturation concentration of peptide effects in the STG (Zhao et al., 2011). In only one of the four cases, co-modulation increased the effect. For the PD to LP synapse, Proc did not completely occlude the effect of adding CCAP, probably because saturating Proc receptors alone does not fully activate the target. In the other three cases, co-application did not produce an additional effect (Figure 4). The fact that complete occlusion was achieved in both synapses by at least one peptide confirms that synapse modulation was maximal when both peptides were applied at 1 μM.

**Figure 4.**
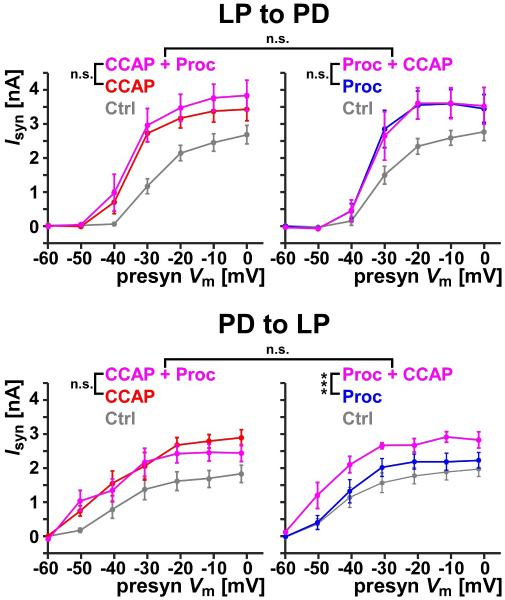
Maximum co-modulation of the synaptic currents by 1 μM CCAP and 1 μM Proc. Each panel shows the effect of co-modulation of either synapse on the synaptic activation curve, following modulation by 1 μM of either modulator alone. For the LP to PD synapse (top), co-modulation did not increase the synaptic current significantly compared to either CCAP alone (left, p=0.45, N=5) or Proc alone (right, p=1.0, N=6). Between the two sets of experiments (left and right panels), neither control levels (p=0.55), nor co-modulation levels (p=0.68) were significantly different. For the PD to LP synapse (bottom), co-modulation did not increase the synaptic current significantly compared to CCAP alone (left, p=0.99, N=6), but it did increase the effect of Proc alone (right, p <0.0001, N=6). Once again, between the two sets of experiments (left and right panels), neither control levels (p=0.73), nor co-modulation levels (p=0.47) were significantly different. All statistical comparisons were Two-Way RM-ANOVA, followed by a Tukey post hoc analysis, if applicable. (* p<0.05, *** p<0.001).

Notably, at both synapses, co-modulatory effects were not dependent on the order of application. Synaptic activation curves were not statistically different between experiments in which either CCAP or Proc were applied first (Figure 4). We also verified that the control measurements were not different for each synapse. Therefore, although co-modulation may have additional effects depending on the neuromodulator and the synapse, the saturation level of synaptic co-modulation was not history dependent.

### Neither CCAP nor Proc modulates short-term synaptic plasticity

The pyloric circuit is rhythmically active with a frequency between ~0.5 and 2 Hz (Goaillard et al., 2009). Like many synapses in the STG, the LP to PD and PD to LP synapses exhibit short-term synaptic depression (Tseng and Nadim, 2010; Zhao et al., 2011). In rhythmically active circuits, short-term synaptic plasticity means that the strength of the synapse depends on the period of the rhythm (Manor and Nadim, 2001). This means that depressing synapses are the stronger the faster the rhythm is, whereas the opposite is true for facilitating synapses. Hence, neuromodulation of short-term synaptic plasticity can play an important role in shaping circuit output and dynamics.

At both synapses, we found that neither CCAP, nor Proc, nor co-application of both, significantly changed the level of short-term synaptic depression with a presynaptic voltage step of 40 mV amplitude (Figure 5). This is consistent with a prior study of the effects of Proc on the PD voltage responses to large LP depolarizations (Zhao et al., 2011). In the same study, Zhao et al. 2011 described a significant effect on short-term synaptic dynamics when smaller presynaptic voltage-steps were used. However, a detailed analysis of the voltage-dependence of modulatory effects on synaptic plasticity exceeded the scope of our study.

**Figure 5.**
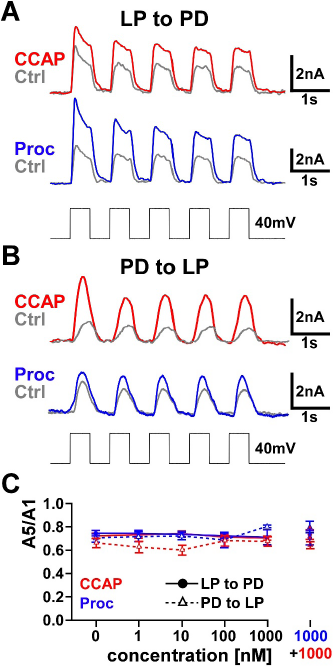
CCAP, Proc or combinations of both do not modulate short-term synaptic plasticity measured with large presynaptic voltage steps. **(A, B)**: Sample experimental traces showing the five postsynaptic currents (with mean amplitude Amp1-Amp5) in response to a set of five presynaptic voltage steps from −60 mV to −20 mV in control and in the presence of either modulator, for the LP to PD (**A**) and PD to LP (**B**) synapses. **(C)** Short-term synaptic plasticity was quantified as Amp5/Amp1. This ratio did not change from control to different concentrations of individual neuromodulators, or co-modulation. (LP to PD: from control to either CCAP or Proc to co-modulation, p=0.50 and 0.34. PD to LP: from control to either CCAP or Proc to co-modulation, p=0.20 and 0.11. N=6 for all. All measurements One-Way RM ANOVA).

### Co-modulatory effects on synapses are linearly additive up to saturation

After establishing that the saturation level of co-modulation is not history dependent, we used equation (6) to calculate the co-modulation predictions for the synapses. Recall that the individual effects of the two peptides were modeled by the predictors for their dose-dependent effects (equation (3) and Figure 3). The linear summation rule predicts that the co-modulatory effect is the sum of the individual modulatory enhancements due to Proc and CCAP at their respective concentrations (equation (4)), up to saturation. We tested this prediction on both synapses with 18 different modulator combinations (see Methods).

We compared our predictions with the experimental results by computing the coefficient of determination (R^2^, evaluating the trend of the data) and normalized root mean squared error (NRMSE, evaluating the deviation of the data from the prediction; see Methods). We report these statistics for each combination individually, and also report the overall R^2^ and NRMSE for all combinations.

For the LP to PD synapse, our prediction matched the experimental results exceedingly well (examples shown in Figure 6A, all data provided in Figure 6-source data). The comparison between predicted and measured values showed high prediction accuracy (Figure 6B, the line *y = x* indicates a perfect match). For all combinations, we obtained high R^2^ and low NRMSE values, indicating that our predictions both captured the trend of the data well and had negligible deviation from the data (Figure 6C; see Figure 6-figure supplement for exact values). The overall values were R^2^ = 0.90 and NRMSE = 0.31 for this synapse. We therefore concluded that co-modulation of LP to PD synapse can be predicted from effects of individual peptides using the linear summation rule.

**Figure 6.**
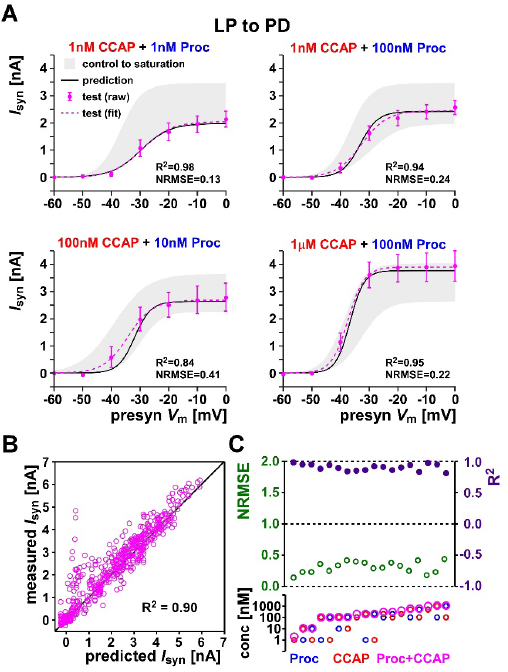
The co-mod ulatory effect of CCAP and Proc on the LP to PD synapse can be predicted from linear summation up to saturation. **(A)** The LP to PD synaptic current activation curve in response to co-applied CCAP and Proc at four different concentration combinations (test, raw and fit) is well predicted by the model (prediction). Also shown is the range of synaptic currents measured in the respective experiments (control to saturation). The R^2^ and NRMSE values in each case show the goodness of the prediction. **(B)** The prediction values compared with the actual measurements for all data points in the 18 different combinations of co-modulation measurements of the LP to PD synapse. Also shown, for comparison, are the line of perfect prediction *(y=x)* and overall R^2^ values. All data points are provided in Figure 6-source data. **(C)** The R^2^ and NRMSE values shown for each of the 18 co-modulation combinations of the LP to PD synapse. R^2^=1 and NRMSE=0 indicate perfect predictions, whereas R^2^=0 and NRMSE=1 indicate that the prediction was no better than the mean of the data. The bottom panel shows the concentration of Proc, CCAP and total concentration (Proc+CCAP) in each case. Data are shown in order of increasing total concentration. Each combination included 5-6 preparations.

We observed simililar accuracy of the linear prediction for the PD to LP synapse (examples shown in Figure 7A, all data provided in Figure 7-source data). The predictions for the PD to LP synapse also had high R^2^ and low NRMSE, with an overall R^2^ = 0.73 and NRMSE = 0.52 (Figure 7B and C; see Figure 7-figure supplement for exact values). These values indicate that co-modulation of the PD to LP synapse was predicted well by the linear summation rule, if not quite as accurately as at the LP to PD synapse.

**Figure 7.**
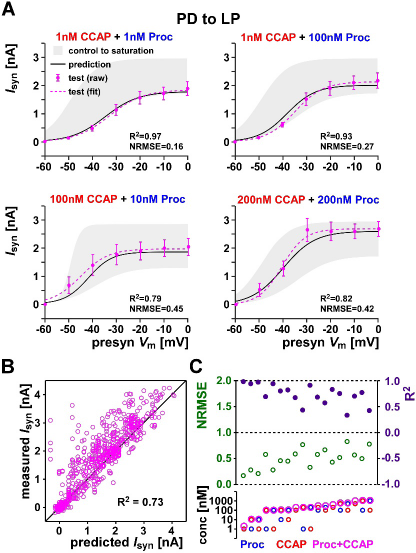
The co-modulatory effect of CCAP and Proc on the PD to LP synapse can be predicted from linear summation up to saturation. **(A)** The PD to LP synaptic current activation curve in response to co-applied CCAP and Proc at four different concentration combinations (test, raw and fit) is well predicted by the model (prediction). Also shown is the range of synaptic currents measured in the respective experiments (control to saturation). The R^2^ and NRMSE values in each case show the goodness of the prediction. **(B)** The prediction values compared with the actual measurements for all data points in the 18 different combinations of co-modulation measurements of the PD to LP synapse. Also shown, for comparison, are the line of perfect prediction *(y=x)* and overall R^2^ values. All data points are provided in Figure 7-source data. **(C)** The R^2^ and NRMSE values shown for each of the 18 co-modulation combinations of the LP to PD synapse. R^2^=1 and NRMSE=0 indicate perfect predictions, whereas R^2^=0 and NRMSE=1 indicate that the prediction was no better than the mean of the data. The bottom panel shows the concentration of Proc, CCAP and total concentration (Proc+CCAP) in each case. Data are shown in order of increasing total concentration. Each combination included 5-6 preparations.

### Co-modulatory effects on *I*_MI_ are not linearly additive

Our data indicate that the co-modulatory effects of Proc and CCAP on the synapses were linearly additive, up to saturation. This suggests that the intracellular pathways underlying the Proc and CCAP effects converge in the LP and PD neurons, without additional interactions. If so, it is reasonable to assume that the activation of *I*_MI_ by Proc and CCAP would also follow the same rule.

The protocols that we used to measure the synaptic current from LP to PD also allowed us to estimate the level of *I*_MI_ in the LP neuron (see Methods and Figure 8A). We therefore quantified the dose-dependent activation of *I*_MI_ in the presence of either Proc or CCAP. Both peptides activated *I*_MI_ starting at nanomolar concentrations and consistently produced larger currents as the concentration increased (Figure 8B and C).

In each experiment, either Proc or CCAP was applied at increasing concentrations up to 1μM, and then both peptides were co-applied at 1 μM each. Co-application revealed complete occlusion in both directions and did not show history dependence (Figure 8C): The addition of the second peptide did not significantly increase the *I*_MI_ response, and *I*_MI_ values were not significantly different between the different orders of application.

The dose-dependent curves for the two peptides were used to construct the predictors of the co-modulation effect (equation (5)). From these individual predictors, we calculated the *I*_MI_ levels expected to be activated by each peptide at any concentration, using linear summation to saturation (equation (7)). As with the synapses, we compared the predicted *I*_MI_ levels to the actual measurements in 18 different co-modulation combinations. We then calculated the R^2^ and NRMSE values for each individual combination and for all combinations together. For these comparisons, *I*_MI_ was measured at −15 mV. Calculations of the R^2^ and NRMSE values with the peak *I*_MI_ level, derived from the fitted IV curves (equation (2)), produced similar results (Figure 8-source data).

Surprisingly, and in stark in contrast to the synapses, our predictions were far from the measured values of the co-modulated *I*_MI_ in the LP neuron (Figure 8E top). The comparison between predicted and measured *I*_MI_ value shows over-estimation in most of the data points (Figure 8D). For half of the combinations, R^2^ values were below 0 and NRMSE values were above 1 (Figure 8E middle; see Figure 8-figure supplement for exact values). The low overall R^2^ value of 0.08 and high overall NRMSE value of 0.96 indicate that our linear summation model was a very poor predictor for the co-modulation of *I*_MI_ and in fact no better than using the mean of the data as a predictor.

**Figure 8.**
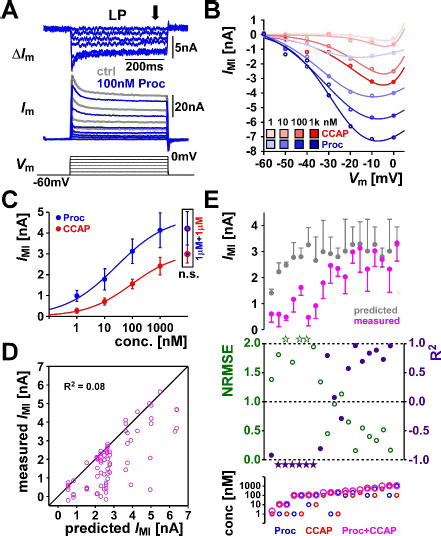
The co-modulatory effect of CCAP and Proc on the levels of *I*_MI_ in the LP neuron cannot be predicted from linear summation up to saturation. **(A)** Measurement of *I*_MI_ in the LP neuron. The total membrane current (*I*_M_) was measured in the LP neuron, with 500 ms voltage steps from −60 to 0 mV, in control saline and in the presence of 100 nM Proc. The difference current (∆/m) was calculated by digital subtraction and *I*_MI_ was calculated as the mean current in the latter half of the voltage step (arrow). **(B)** Example of the *I*_MI_ IV curves measured in two experiments in increasing concentrations of CCAP or Proc, shown together with the fit of the data points using equation (2). **(C)** Dose-dependent levels of *I*_MI_ (measured at −15 mV) in the presence of Proc, CCAP or both (1 μM of each). Dose-dependent parameters, for CCAP: *I*_max_ = 3.080, *C*_mid_= −7.025, *C*_c_= 0.7997; for Proc: *I*_max_ =4.797, *C*_mid_= −7.568, *C*_c_= 0.8699. *I*_MI_ measured with co-applied 1 μM CCAP and Proc in the two sets of the experiments were not significantly different (Student's t-test, p = 0.31, N=6 for both sets of experiments.) **(D)** The linear-summation-up-to-saturation prediction values compared with the actual measurements of *I*_MI_ for all data points in the 18 different combinations of co-modulation. Also shown, for comparison, are the line of perfect prediction *(y=x)* and overall R^2^ values. All data points are provided in Figure 8-source data. **(E)** Measured (at −15 mV) and predicted *I*_MI_ values, as well as the R^2^ and NRMSE values shown for each of the 18 co-modulation combinations. R^2^=1 and NRMSE=0 indicate perfect predictions, whereas R^2^=0 and NRMSE=1 indicate that the prediction was no better than the mean of the data. Stars indicate out of range values. The bottom panel shows the concentration of Proc, CCAP and total concentration (Proc+CCAP) in each case. Data are shown in order of increasing total concentration. Each combination included 4-5 preparations.

Interestingly, also in contrast to the fairly consistent R^2^ and NRMSE values across different co-modulation combinations for the synapses, these values varied drastically across different combinations for *I*_MI_ (Figure 8E middle). The predictor did very poorly (NRMSE>1) when at least one of the peptide was at a low concentration, but somewhat better (NRMSE closer to 0) when the combined concentrations were high, mostly because the predictor estimated the co-modulation to be at saturation (Figure 8E).

Despite the poor prediction, our linear model provided some useful information about the dynamics of *I*_MI_ co-modulation. The measured *I*_MI_ level was always lower than the prediction, indicating that the co-modulatory effect was sublinear.

## Discussion

### Distinct rules for co-modulation of different subcellular targets

It is common for multiple neuromodulators to target the same ion channel or synapse, or have distinct targets within the same neuron (McCormick and Williamson, 1989; Harris-Warrick, 2011; Marder, 2012). As such, circuit output depends on how signaling pathways mediated by distinct neuromodulator receptors converge and interact. The actions of converging neuromodulators may have the same or opposing signs (Nadim and Bucher, 2014). Regardless of the signs of the action, converging neuromodulators could have additive, synergistic, antagonistic, or other nonlinear co-modulatory effects. For a given target, it is important to know if convergent neuromodulators act in a simple additive manner or have more complex, nonlinear interactions. An additional open question is whether the interactions of neuromodulators that converge onto multiple subcellular targets follow the same rule at all shared targets.

Despite recent advances in genetic and imaging tools (Arrigoni and Saper, 2014; Cohn et al., 2015; Shahidi et al., 2015), many systems still lack experimental accessibility or the basic understanding of neuromodulator actions on their cellular and subcellular targets to explore this topic. Peptide neuromodulation of the pyloric circuit of the STG provides a special opportunity to explore the rules of co-modulation of synaptic and intrinsic ionic currents, and to understand their consequences at the circuit level (Daur et al., 2016). We observed linearly additive co-modulation of synapses, but sub-linearly additive co-modulation of a voltage-gated ionic current in the same neurons. These specific results may be idiosyncratic for the neurons and synapses we studied, as co-modulation of synapses can be nonlinear (Parker, 2000), and co-modulation of voltage-gated ionic currents could be linearly additive. However, the important lesson from our findings is that converging co-modulation of synapses and ionic currents by the same neurmodulators, or different subcellular targets in general, can follow distinct rules. Given the complex patterns of divergence and convergence of neuromodulators in many systems, this finding likely has broad functional implications.

### Linearly additive co-modulation of pyloric synapses

Modulation of synaptic currents may involve both presynaptic changes in transmitter release and postsynaptic changes in ionotropic receptor properties. Therefore, the total effect can result from modulation of molecular components in two different neurons, involving potentially distinct signaling pathways and concentration dependence. At synapses in the STG, a single neuromodulator can exert functionally opposing effects on the pre- and postsynaptic sides, for example enhancing transmitter release but reducing postsynaptic responsiveness (Harris-Warrick and Johnson, 2010; Garcia et al., 2015). We therefore did not necessarily expect co-modulation of synapses to be simply linearly additive. Surprisingly, we observed such linearly additive co-modulation at both synapses. For the LP to PD synapse, CCAP modulation must be presynaptic, as PD neurons do not express CCAP receptors (Garcia et al., 2015). However, Proc modulation could have both pre- and postsynaptic components. Although Proc receptor expression in these neurons has not been tested because their molecular identity has not been determined in the STG, both neurons show *I*_MI_ activation in response to Proc application (Swensen and Marder, 2000). For the PD to LP synapse, both modulators could have pre- and postsynaptic effects. The synaptic input to the LP neuron from the pacemaker (which we measured as the PD to LP synapse) is from both AB and PD neurons. AB expresses CCAP receptors (Garcia et al., 2015) and isolated AB neurons respond to both CCAP and Proc (Swensen and Marder, 2001). The fact that we measured the synaptic responses of LP while voltage clamping only one of the presynaptic neurons may explain why linear summation less accurately predicts co-modulation compared to the LP to PD synapse (Figures 6 and 7).

We did not investigate whether neuromodulatory effects occurred pre- or postsynaptically, or both. However, given that we observed linear summation and occlusion, it is likely that modulatory signaling on either side was purely converging, without any nonlinear interactions. Linear co-modulation could also occur through spatial segregation, for example, when one neuromodulator only acts presynaptically, and the other only postsynaptically. Even in a single neuron, modulatory micro-domains can provide non-overlapping, independent activation of identical targets using the same signaling pathways (Lur and Higley, 2015). However, in the case of spatial segregation, no occlusion should occur, and the saturation level of co-modulation should be the linear sum of the maximum effects achieved by each neuromodulator.

### Sublinear co-modulation of *I*_MI_

In contrast to the synapses, we observed nonlinear co-modulation of *I*_MI_, which Indicated that the signaling pathways targeting *I*_MI_ were distinct from the pathways targeting the synapses. It was previously suggested that peptides modulate synapses in the STG through their actions on the *I*_MI_ channel, which might be partially permeable to calcium (Zhao et al., 2011; Gray et al., 2017). However, our results indicate that this is unlikely, given that linear co-modulation of the synapses and nonlinear co-modulation of *I*_MI_ occurred in the same experiments. The nonlinearity of *I*_MI_ co-modulation may have two components: sublinear interactions when at least one modulator is at low concentration, and occlusion when both are at high concentrations (Figure 8E). The occlusion effect was also shown in our previous study (Garcia et al., 2015).

In *C. borealis,* the Proc receptor gene has not been identified, and there appears to be only one CCAP receptor gene (Garcia et al., 2015). In insects, Proc receptors come from a single gene (Caers et al., 2012). Different CCAP receptor genes have been found to produce receptors that differ more than 30-fold in their agonist affinities (Li et al., 2011), but the underlying gene duplication is thought to have occurred only in some insect lineages. However, this does not exclude the possibility of post-translational modifications that could result in receptors with different agonist affinities or differential activation of different signaling pathways (Leclerc et al., 2006; Daaka, 2012). This opens the possibility that in the STG, peptides activate *I*_MI_ through receptor subtypes with different affinities. If so, the low- and high-affinity pathways mediated by the same peptides should undergo simple convergence, because the dose-dependent activation of *I*_MI_ is sigmoidal (Figure 8C). Similarly, the low-affinity receptor mediated pathways should also converge without lateral interactions, resulting in occlusion at high concentrations. However, the high-affinity pathway mediated by one peptide might inhibit the low-affinity pathway mediated by the other, possibly by targeting the intracellular calcium concentration or calcium-binding proteins (Gray et al., 2017), thus reducing the *I*_MI_ level activated by the low-affinity pathway. Such an interaction may remain distinct from the linear additive rule of the LP to PD synapses, e.g., if the synaptic neuromodulation pathway is through distinct signaling molecules activated by these receptors.

Another possible mechanism is that the CCAP and Proc receptors can form a heteromer complex and display behaviors distinct from either receptor alone (reviewed in Smith and Milligan, 2010). Given the variety of possible mechanisms, a different set of experiments, as well as mathematical modeling, will be required to provide an accurate description of the co-modulation rule for *I*_MI_.

### Distinct co-modulation rules may increase flexibility and functionally uncouple the modulation of different targets

When different neuromodulators converge onto multiple targets, their actions on the shared targets are inextricably linked. However, modulator effects on different targets can be uncoupled by different co-modulation rules. For example, in the results shown here, 1 nM CCAP and 100 nM Proc produced an additive effect in the LP to PD synapse, but activated much less *I*_MI_ than 100 nM Proc alone (Figures 6A, 8C and 8E). In the pyloric circuit, *I*_MI_ enhances neuronal excitability of the pacemaker neurons and thereby regulates the pyloric frequency (Hooper and Marder, 1987). The synapses from the pacemaker neurons (AB and PD) to follower neurons like LP are important for the regulation of burst phasing across pyloric neurons (Eisen and Marder, 1984; Rabbah and Nadim, 2005; Goaillard et al., 2009). The feedback synapse from LP to PD has little effect on the mean rhythm frequency, but reduces its variability (Zhao et al., 2011).

Distinct rules for co-modulation of neuronal excitability and synaptic interactions could functionally uncouple these effects and therefore allow burst phasing and rhythm frequency to be regulated differentially.

Furthermore, sub-linear co-modulation of *I*_MI_ may extend the dynamic range for the modulation of neural excitability by producing qualitatively different effects than each individual neuromodulator. Because STG neurons are modulated by many peptides, sublinear co-modulation would ensure that neuronal excitability is not saturated during baseline activity when many peptides may be present at low concentrations. Yet, when any specific peptide neuromodulator is released at a higher concentration, it can produce a distinct circuit output.

### Co-modulation in light of animal-to-animal variability

Across individuals, pyloric neurons display substantial variability in the magnitude of synaptic and voltage-gated ionic currents, as well as in the expression levels of mRNAs that code for ion channels (Golowasch et al., 1999; Schulz et al., 2006; Schulz et al., 2007; Goaillard et al., 2009). Despite this variability, which is several-fold in some cases, neuronal excitability and the patterning of circuit activity is well maintained (Bucher et al., 2005; Goaillard et al., 2009; Marder et al., 2015). Substantial variability has also been described for neuromodulatory components. For example, CCAP receptor mRNA expression varies 3-fold in the LP neuron (n=22 in Garcia et al., 2015), and CCAP-activated *I*_MI_ in the LP neuron varies more than 5-fold in amplitude (n=15 in Goaillard et al., 2009). There may also be long-term regulatory changes in neuromodulation, perhaps due to seasonal or molt cycle related hormonal changes, which are almost impossible to control for in wild caught animals. In contrast to the data presented here, in a previous study we found that, in LP, CCAP activated a larger *I*_MI_ than Proc did, and the Proc response was not saturating (Garcia et al., 2015). In this study, we only tested each co-modulation combination on a small number of animals (n=4-6), but the total number of animals we used in this study (n=33) matched the variability of *I*_MI_ levels seen in the previous studies. The fact that, despite this variability, the linear summation rule accurately predicted co-modulation of the synapses indicates that, co-modulation rules appear to be robust across individuals, despite component variability.

### Bridging levels of co-modulation effects

Unraveling the consequences of co-modulation at the circuit level requires examining their interactions at multiple levels. In this study, we took a first step toward identifying the rules of co-modulation at the level of shared targets. However, our study leaves several questions unanswered.

First, the signaling pathways resulting in our observed data remain unknown. Second, we bath applied neuromodulators in our study, which was necessary to quantify precise dose-dependent effects, but as a number of studies in the STG have shown, fails to address the spatiotemporal dynamics of neuromodulation (Nusbaum et al., 2017). Neuromodulators can be released as hormones or as neurotransmitters. In the latter case, spatiotemporal properties of synaptic transmission can be critical in determining circuit output (reviewed in Nusbaum et al., 2017). The spatial interactions depend on the architecture of the local circuits, the spatial pattern of neuromodulator release and the peptidase activity. For neurotransmitter modulators, the temporal dynamics is, by necessity, determined by the patterns of activity of the modulatory neurons that release these transmitters. The activity patterns of the modulatory neurons, in turn, is subject to feedback from the activity of the target circuits, thereby producing another potential level of complexity. To probe the spatiotemporal dynamics of co-modulation, combining experimental approaches, such as stimulating neuromodulatory projection neurons, and computational modeling is necessary.

Finally, all our experiments were done with voltage-clamp steps in order to characterize the neuromodulatory effects on each target. However, such experiments mask the interactions among circuit components, both those within neurons and those with their synaptic partners. One such example is shown in (Zhao et al., 2011) for the LP to PD synapse, where Proc changes two factors: it enhances both the burst voltage waveform of the presynaptic LP neuron and the amplitude of the synaptic current. When the LP neuron is voltage clamped with the pre-recorded realistic control or Proc voltage waveforms, the resulting synaptic currents are similar in control saline, but different in the presence of Proc. This indicates that the first factor (change in the LP waveform) produces a meaningful effect only in conjunction with the second factor (direct enhancement of synaptic release). Exploring such interactions among cellular or circuit components is important in understanding the functional consequences of co-modulation and requires further experiments and computational modeling.

## Conclusions

The persistent actions of neuromodulators are critical for proper circuit function and plasticity. Because neuromodulators do not act independently, understanding their interactions at different concentrations is fundamentally important for the understanding of circuit dynamics and resulting behaviors. Identifying the mechanisms of co-modulation also provides mechanistic guidance for therapies that target one or more neuromodulatory pathways (Engineer et al., 2011; Pena et al., 2014; Freret et al., 2017). Here, we made a first step towards the goal of understanding how neuromodulators interact to shape the circuit output, by quantitatively clarifying the co-modulatory rules at target level. Given co-modulation is a universal and evolutionarily conserved strategy, our results can provide insights and new hypothesis to test at system level. We also provide an initial framework to test similar rules in other circuit components, other neuromodulators and other systems. However, the challenge will remain to translate findings from the level of ionic currents to the effects of co-modulation on actual synaptic function and neuronal excitability, and from there to circuit activity. Even in small circuits with identified neurons, as the pyloric circuit used here, this will require a multipronged approach, combining multiple experimental and computational methods (Nadim and Bucher, 2014).

## Materials and Methods

### Preparation and electrophysiological recordings

All experiments were done on wild-caught adult male crabs *(Cancer borealis)* purchased from local seafood stores. Prior to experiments, animals were kept in artificial sea water tanks at 13 °C. Before dissection, crabs were anesthetized by placing on ice for at least 30 min. The STNS was dissected out following standard protocols (Blitz et al., 2004; Tohidi and Nadim,2009), placed in a Petri dish coated with clear silicon elastomer (Sylgard 184; Dow Corning; Midland, MI) and superfused with *C. borealis* saline, containing (in mM) 11 KCl, 440 NaCl, 13 CaCl2, 26 MgCl2, 11.2 Trizma base, and 5.1 maleic acid (pH =7.4 −7.5). A petroleum jelly well was built around the STG for constant superfusion of chilled (10-12 °C) saline during the experiment.

For neuron identification, extracellular motor nerve recordings were obtained with a differential AC amplifier (A-M Systems, Model 1700; Sequim, WA), using stainless-steel pin wire electrodes placed inside and outside of small petroleum jelly wells built around the nerves. Intracellular recordings and voltage clamp were done with Axoclamp 900A amplifiers (Molecular Devices; San Jose, CA). The STG was desheathed and the neuron somata were impaled with sharp glass electrodes, pulled with a Flaming-Brown P-97 Puller (Sutter Instruments; Novato, CA) and filled with 0.6 M K2SO4 + 20 mM KCl solution (15-30 MΩ electrode resistance). Neurons were identified by their characteristic intracellular waveforms and by matching their activities to the spikes on the corresponding motor nerves. All electrophysiological data were digitized at 5-10 KHz with a Digidata 1440A data acquisition board (Molecular Devices).

### Neuromodulatory effects on the strength and dynamics of the synaptic currents

The neuromodulatory effects on strength and short-term plasticity of the graded component of both the LP to PD and the PD to LP synapses were measured with simultaneous dual two-electrode voltage clamp recordings of the PD and LP neurons.

In voltage clamp experiments, 10 nM tetrodotoxin citrate (TTX; Biotium; Fremont, CA) saline was bath applied to block action potentials and descending neuromodulatory inputs. The synaptic current was measured as the current elicited in the postsynaptic neuron (held at −50 mV), in response to depolarizing 500-1000 ms voltage steps in the presynaptic neuron (from a holding potential of −60 mV to 0 mV, in 10 mV steps; Figure 1 B and C). The postsynaptic current reported in this study is the mean value of the current during the first 500 ms of the presynaptic pulse (the postsynaptic current integral divided by the presynaptic voltage step duration of 500 ms). The peak values of the synaptic currents during each voltage step are included in Figure 2-source data.

To fit the postsynaptic current amplitude as a function of presynaptic voltage *(V_pre_),* we used a sigmoid function of the following form:

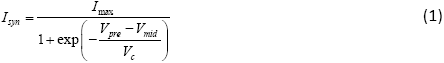

In these fits, we assumed that the postsynaptic current was 0 at *V_pre_* = −70 mV.

Proc (Bachem; Torrance, CA and Genscript; Piscataway, NJ) and CCAP (Bachem) were aliquoted in 1 mM stock solutions and stored at −20 °C until use. For each experiment, the aliquots were further diluted to the desired concentrations. The dose-dependent effect of Proc or CCAP on synapses was measured by bath applying each peptide from low to high concentration (1 nM to 1 μM) with a four-minute interval between each concentration. We considered 1 μM to be the saturation concentration of both Proc and CCAP based on previous studies (Zhao et al., 2011). In addition, 1μ M Proc and CCAP were co-applied at the end of each experiment to measure the maximum modulatory effect.

To measure short-term synaptic plasticity, we voltage clamped the presynaptic neuron at a holding potential of −60 mV and applied a set of five 500 ms identical depolarizing square pulses, from −60 to −20mV, at 1Hz. We measured the mean current amplitude in the postsynaptic neuron (voltage clamped at −50 mV) in response to each pulse. The level of shortterm plasticity was quantified as the ratio of the postsynaptic current amplitude elicited by the fifth and first pulses. For the experiments that had two repeated measurements, we averaged the two measurements.

### Neuromodulatory effects on the voltage-gated ionic current *I*_MI_

The modulator-activated inward current *I*_MI_ was measured in the LP neuron in the same experiments in which we measured the LP to PD synaptic current. Because, in these experiments, the LP neuron membrane potential was stepped from −60 to 0 mV for measuring the LP to PD synapse (using the current measured in the postsynaptic PD neuron), the same voltage steps could be used to estimate *I*_MI_ in the LP neuron (using the voltage-clamp current, *I*_LP_, injected in the presynaptic LP neuron). *I*_MI_ was measured as the difference between *I_LP_* measured in the presence of the modulator and *I_LP_* measured in control saline (Figure 8A) (Golowasch and Marder, 1992). *I*_MI_ is a non-inactivating current (Golowasch and Marder, 1992; Gray et al., 2017). To reduce errors due to differences in transient currents, we reported the mean value of the difference current, measured in the second half of each voltage pulse where the currents had reached approximate steady state. The *I*_MI_ value at −15 mV was measured as the average of the currents elicited at −20 mV and −10 mV step voltage and used for analysis.

*I*_MI_ is a non-inactivating fast voltage-gated inward current whose activation curve is a simple Boltzmann sigmoidal equation (Goaillard et al., 2009). The IV curve of *I*_MI_ can therefore be estimated as

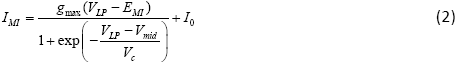

Where *g_max_* is the maximum conductance of *I*_MI_, **E*_MI_* is the reversal potential and *I*0 is the baseline difference current.

The dose-dependent effects of the modulators and the protocols for co-modulation of *I*_MI_ were the same as those described for the synapses above.

### Constructing predictors for single neuromodulators

For each neuromodulator-synapse pair, we fit a surface to the postsynaptic currents measured at all presynaptic voltages and concentrations in multiple experiments. The equation used to define this surface was a dual sigmoidal function of both the presynaptic voltage *(V_pre_)* and the log peptide concentration (C). This equation was based on equation (1), so that

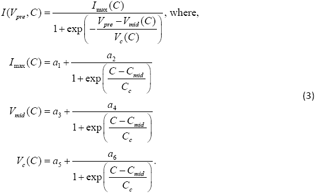

In these fits, the unit of peptide concentration is M, and the control value was set at *C =* −10, thus assuming that 10^−10^ M concentration had no effect. The enhancement functions for each peptide were defined as the increase produced by the modulator above the control level of the synaptic current at each presynaptic voltage:

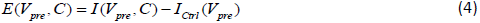

The resulting enhancement functions served as predictors for the effect of the neuromodulator on the postsynaptic current at any voltage and concentration.

In the case of *I*_MI_, we fit the dose-dependent effects of Proc and CCAP with the sigmoidal curve

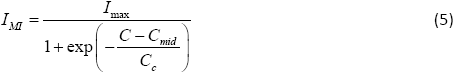

where *C* is the log peptide concentration and *C*_mid_ and *C*_c_ are, respectively, the half-maximum log concentration and the slope factor. In these fits, the unit of peptide concentration is M, and the control value was set at *C* = −10, assuming that 10^−10^ M concentration had no effect.

### Predicting and testing co-modulation

We compared the predictions of co-modulation effects with the experimental data from co-applications of Proc and CCAP in 18 different combinations of concentrations for both the LP to PD and PD to LP synapses and *I*_MI_ in the LP neuron. These 18 combinations were divided into four separate groups of experiments, with each group only containing four or five combinations (group information can be found in Figure 6 - Figure supplement). In each group of experiments,each peptide was applied in order from lower to higher concentration. Each combination was bath applied for a four-minute-interval, a value calculated by the superfusion rate, the volume of solution in the line and the size of the petroleum jelly well around the STG. At the end of each experiment, Proc and CCAP were co-applied at 1 μM each to record the maximum modulatory effect in that preparation.

The predictions for synapses were calculated by adding up the enhancements produced by each peptide at the respective concentrations (obtained from equation (4)) and the control value (*I_ctrl_co_mod_*), and limiting the sum to the saturation level ( *I_sat_co_mod_*) which is the synaptic current elicited by both peptides co-applied at 1μM.

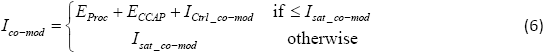

For each combination, we measured the co-modulated synaptic currents, as described above, at presynaptic voltages from −60 mV to 0 mV, in 10 mV steps. We then compared the measurement with the prediction for those voltages.

The co-modulation predictions for *I*_MI_ were calculated by simply adding up the value of *I*_MI_ activated by each modulator at its respective concentration on the dose-response curve, limited to the saturation level.

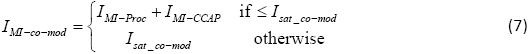

To estimate how well our model prediction fit the experimental results, we used two standard goodness-of-fit tests. One of these measures is the coefficient of determination R^2^ measured as: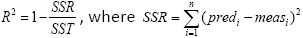 is the summed square of the residuals and 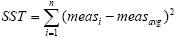 is the total sum of squares. R^2^ − 1 means that the prediction perfectly captures the trend of the data. Note, however, that this R^2^ is different from the Pearson correlation coefficient where a linear fit to the data is evaluated. In our case, R^2^ may be < 0, which simply indicates that the mean of the data *meas_avg_* provides a better prediction than the model.

The second measure we use is the normalized root mean squared error (NRMSE,normalized to standard deviation), calculated as 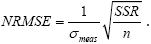 A value of 0 for NRMSE indicates a perfect fit, whereas values > 1 indicate that the mean of the data *meas_avg_* provides a better prediction than the model.

We report both R^2^ and NRMSE as recommended by Schunn and Wallach (Schunn and Wallach, 2005) to show that our prediction captures both the trend of the data and how far it deviates from the exact data points. Specifically, R^2^ evaluates whether the model prediction captures the trend of the data, whereas, NRMSE evaluates the deviation of the data from the prediction.

### Data analysis and statistical analysis

All data and statistical analysis were done with Matlab (MathWorks, 2015b; Natick, MA) and R (The R Foundation). Unless otherwise indicated, all error bars represent standard error of the mean. Statistical tests included Student’s t-test, One- or Two-way RM ANOVA (followed by post hoc pairwise comparisons done with the Tukey method, when applicable). Critical significance level was set to α=0.05. Comparisons between model prediction and the data were done by reporting the adjusted R^2^ and normalized root mean square error (NRMSE) analysis, as described above.

---

**Figure 6 - Figure supplement.**
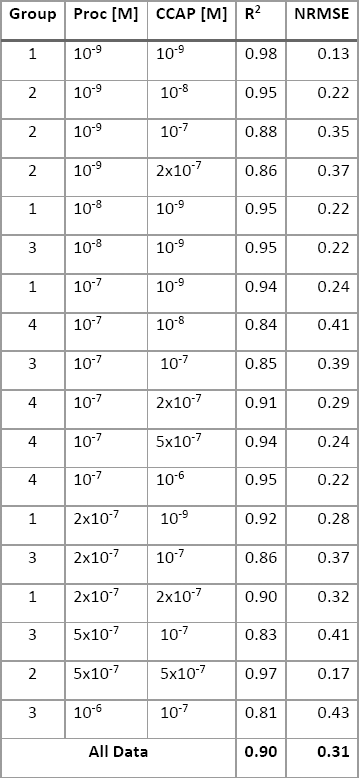
Statistics of the LP to PD synapse co-modulation. Co-applications with the same Group number were performed in the same experiments.

**Figure 7 - Figure supplement.**
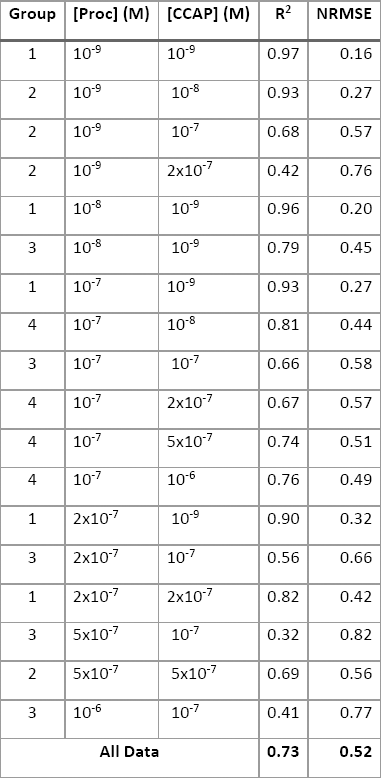
Statistics of the PD to LP synapse co-modulation. Co-applications with the same Group number were performed in the same experiments.

**Figure 8 - Figure supplement.**
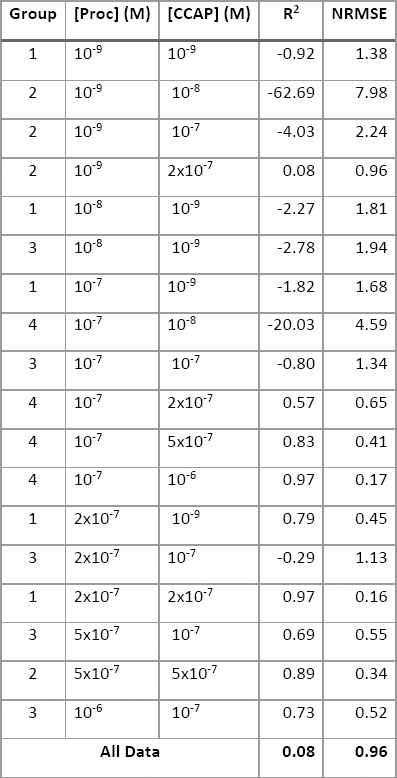
Statistics of the *I*_MI_ co-modulation. Co-applications with the same Group number were performed in the same experiments.

## Acknowledgements

This work was supported by NIH Grants MH060605 and NS083319.

## References

Arrigoni E, Saper CB (2014) What optogenetic Stimulation is telling us (and failing to tell us) about fast neurotransmitters and neuromodulators in brain circuits for wake-sleep regulation. Curr Opin Neurobiol 29:165–171.

Asahina K (2017) Neuromodulation and Strategic Action Choice in Drosophila Aggression. Annu Rev Neurosci 40:51–75.

Bargmann CI (2012) Beyond the connectome: how neuromodulators shape neural circuits. Bioessays 34:458–465.

Beliez L, Barriere G, Bertrand SS, Cazalets JR (2014) Multiple monoaminergic modulation of posturo-locomotor network activity in the newborn rat spinal cord. Front Neural Circuits 8:99.

Blitz DM, Beenhakker MP, Nusbaum MP (2004) Different sensory systems share projection neurons but elicit distinct motor patterns. J Neurosci 24:11381–11390.

Brezina V (2010) Beyond the wiring diagram: signalling through complex neuromodulator networks. Philos Trans R Soc Lond B Biol Sci 365:2363–2374.

Brezina V, Orekhova IV, Weiss KR (1996) Functional uncoupling of linked neurotransmitter effects by combinatorial convergence. Science (New York, NY) 273:806–810.

Bucher D, Prinz AA, Marder E (2005) Animal-to-animal variability in motor pattern production in adults and during growth. J Neurosci 25:1611–1619.

Burke PG, Kanbar R, Basting TM, Hodges WM, Viar KE, Stornetta RL, Guyenet PG (2015) State-dependent control of breathing by the retrotrapezoid nucleus. J Physiol 593:2909–2926.

Caers J, Verlinden H, Zels S, Vandersmissen HP, Vuerinckx K, Schoofs L (2012) More than two decades of research on insect neuropeptide GPCRs: an overview. Frontiers in endocrinology 3:151.

Cohn R, Morantte I, Ruta V (2015) Coordinated and Compartmentalized Neuromodulation Shapes Sensory Processing in Drosophila. Cell 163:1742–1755.

Daaka Y (2012) S-nitrosylation-regulated GPCR signaling. Biochimica et biophysica acta 1820:743–751.

Daur N, Nadim F, Bucher D (2016) The complexity of small circuits: the stomatogastric nervous system. Curr Opin Neurobiol 41:1–7.

Dickinson PS, Fairfield WP, Hetling JR, Hauptman J (1997) Neurotransmitter interactions in the stomatogastric system of the spiny lobster: one peptide alters the response of a central pattern generator to a second peptide. J Neurophysiol 77:599–610.

Djokaj S, Cooper RL, Rathmayer W (2001) Presynaptic effects of octopamine, serotonin, and cocktails of the two modulators on neuromuscular transmission in crustaceans. J Comp Physiol A 187:145–154.

Doi A,Ramirez JM (2008) Neuromodulation and the orchestration of the respiratory rhythm. Respir Physiol Neurobiol 164:96–104.

Donlea JM, Alam MN, Szymusiak R (2017) Neuronal substrates of sleep homeostasis; lessons from flies, rats and mice. Curr Opin Neurobiol 44:228–235.

Eisen JS, Marder E (1984) A mechanism for production of phase shifts in a pattern generator. J. Neurophysiol 51:1375–1393.

Engineer ND, Riley JR, Seale JD, Vrana WA, Shetake JA, Sudanagunta SP, Borland MS, Kilgard MP (2011) Reversing pathological neural activity using targeted plasticity. Nature 470:101–104.

Filosa A, Barker AJ, Dal Maschio M, Baier H (2016) Feeding State Modulates Behavioral Choice and Processing of Prey Stimuli in the Zebrafish Tectum. Neuron 90:596–608.

Freret T, Lelong-Boulouard V, Lecouflet P, Hamidouche K, Dauphin F, Boulouard M (2017) Co-modulation of an allosteric modulator of nicotinic receptor-cholinesterase inhibitor (galantamine) and a 5-HT4 receptor agonist (RS-67333): effect on scopolamine-induced memory deficit in the mouse. Psychopharmacology (Berl) 234:2365–2374.

Garcia VJ, Daur N, Temporal S, Schulz DJ, Bucher D (2015) Neuropeptide Receptor Transcript Expression Levels and Magnitude of Ionic Current Responses Show Cell Type-Specific Differences in a Small Motor Circuit. Journal of Neuroscience 35:6786–6800.

Goaillard JM, Taylor AL, Schulz DJ, Marder E (2009) Functional consequences of animal-to-animal variation in circuit parameters. Nat Neurosci 12:1424–1430.

Golowasch J, Marder E (1992) Ionic currents of the lateral pyloric neuron of the stomatogastric ganglion of the crab. J Neurophysiol 67:318–331.

Golowasch J, Abbott LF, Marder E (1999) Activity-dependent regulation of potassium currents in an identified neuron of the stomatogastric ganglion of the crab Cancer borealis. J Neurosci 19:RC33.

Gray M, Daudelin DH, Golowasch J (2017) Activation mechanism of a neuromodulator-gated pacemaker ionic current. J Neurophysiol 118:595–609.

Harris-Warrick RM (2011) Neuromodulation and flexibility in Central Pattern Generator networks. Curr Opin Neurobiol 21:685–692.

Harris-Warrick RM, Johnson BR (2010) Checks and balances in neuromodulation. Front Behav Neurosci 4.

He K, Huertas M, Hong SZ, Tie X, Hell JW, Shouval H, Kirkwood A (2015) Distinct Eligibility Traces for LTP and LTD in Cortical Synapses. Neuron 88:528–538.

Hooper SL, Marder E (1987) Modulation of the lobster pyloric rhythm by the peptide proctolin. J Neurosci 7:2097–2112.

Inagaki HK, Panse KM, Anderson DJ (2014) Independent, reciprocal neuromodulatory control of sweet and bitter taste sensitivity during starvation in Drosophila. Neuron 84:806–820.

Leclerc PC, Lanctot PM, Auger-Messier M, Escher E, Leduc R, Guillemette G (2006) S-nitrosylation of cysteine 289 of the AT1 receptor decreases its binding affinity for angiotensin II. British journal of pharmacology 148:306–313.

Li B, Beeman RW, Park Y (2011) Functions of duplicated genes encoding CCAP receptors in the red flour beetle, Tribolium castaneum. J Insect Physiol 57:1190–1197.

Lovett-Barron M, Andalman AS, Allen WE, Vesuna S, Kauvar I, Burns VM, Deisseroth K (2017) Ancestral Circuits for the Coordinated Modulation of Brain State. Cell 171:1411–1423 e1417.

Lur G, Higley MJ (2015) Glutamate Receptor Modulation Is Restricted to Synaptic Microdomains. Cell reports 12:326–334.

Manor Y, Nadim F (2001) Synaptic depression mediates bistability in neuronal networks with recurrent inhibitory connectivity. J Neurosci 21:9460–9470.

Marder E (2012) Neuromodulation of neuronal circuits: back to the future. Neuron 76:1–11.

Marder E, Bucher D (2007) Understanding circuit dynamics using the stomatogastric nervous system of lobsters and crabs. Annual review of physiology 69:291–316.

Marder E, Goeritz ML, Otopalik AG (2015) Robust circuit rhythms in small circuits arise from variable circuit components and mechanisms. Curr Opin Neurobiol 31:156–163.

McCormick DA, Williamson A (1989) Convergence and divergence of neurotransmitter action ir human cerebral cortex. Proc Natl Acad Sci U S A 86:8098–8102.

McCormick DA, Pape HC (1990) Noradrenergic and serotonergic modulation of a hyperpolarization-activated cation current in thalamic relay neurones. J Physiol 431:319–342.

Mena W, Diegelmann S, Wegener C, Ewer J (2016) Stereotyped responses of Drosophila peptidergic neuronal ensemble depend on downstream neuromodulators. Elife 5:e19686.

Mesce KA, Crisp KM, Gilchrist LS (2001) Mixtures of octopamine and serotonin have nonadditive effects on the CNS of the medicinal leech. J Neurophysiol 85:2039–2046.

Mirabeau O, Joly JS (2013) Molecular evolution of peptidergic signaling systems in bilaterians. Proc Natl Acad Sci U S A 110:E2028–2037.

Nadim F, Bucher D (2014) Neuromodulation of neurons and synapses. Curr Opin Neurobiol 29:48–56.

Nusbaum MP, Blitz DM, Marder E (2017) Functional consequences of neuropeptide and small-molecule co-transmission. Nat Rev Neurosci 18:389–403.

Park JY, Spruston N (2012) Synergistic actions of metabotropic acetylcholine and glutamate receptors on the excitability of hippocampal CA1 pyramidal neurons. J Neurosci 32:6081–6091.

Parker D (2000) Presynaptic and interactive peptidergic modulation of reticulospinal synaptic inputs in the lamprey. J Neurophysiol 83:2497–2507.

Pena DF, Childs JE, Willett S, Vital A, McIntyre CK, Kroener S (2014) Vagus nerve stimulation enhances extinction of conditioned fear and modulates plasticity in the pathway from the ventromedial prefrontal cortex to the amygdala. Front Behav Neurosci 8:327.

Rabbah P, Nadim F (2005) Synaptic dynamics do not determine proper phase of activity in a central pattern generator. J Neurosci 25:11269–11278.

Richter C, Woods IG, Schier AF (2014) Neuropeptidergic control of sleep and wakefulness. Anni Rev Neurosci 37:503–531.

Schulz DJ, Goaillard JM, Marder E (2006) Variable channel expression in identified single and electrically coupled neurons in different animals. Nat Neurosci 9:356–362.

Schulz DJ, Goaillard JM, Marder EE (2007) Quantitative expression profiling of identified neurons reveals cell-specific constraints on highly variable levels of gene expression. Proc Natl Acad Sci U S A 104:13187–13191.

Schunn CD, Wallach D (2005) Evaluating Goodness-of-Fit in Comparison of Models to Data. In: Psychologie der Kognition Reden und Vorträge anlässlich der Emeritierung von Werner H. Tack: Saarbrücken Univ.-Präsidentin.

Shahidi R, Williams EA, Conzelmann M, Asadulina A, Veraszto C, Jasek S, Bezares-Calderon LA, Jekely G (2015) A serial multiplex immunogold labeling method for identifying peptidergic neurons in connectomes. Elife 4:e11147.

Smith NJ, Milligan G (2010) Allostery at G Protein-Coupled Receptor Homo- and Heteromers: Uncharted Pharmacological Landscapes. Pharmacological Reviews 62:701–725.

Su J, Sandor K, Skold K, Hokfelt T, Svensson CI, Kultima K (2014) Identification and quantification of neuropeptides in naive mouse spinal cord using mass spectrometry reveals [des-Ser1]-cerebellin as a novel modulator of nociception. J Neurochem 130:199–214.

Svensson E, Grillner S, Parker D (2001) Gating and braking of short- and long-term modulatory effects by interactions between colocalized neuromodulators. Journal of Neuroscience 21:5984–5992.

Swensen AM, Marder E (2000) Multiple peptides converge to activate the same voltage-dependent current in a central pattern-generating circuit. J Neurosci 20:6752–6759.

Swensen AM, Marder E (2001) Modulators with convergent cellular actions elicit distinct circuit outputs. J Neurosci 21:4050–4058.

Taghert PH, Nitabach MN (2012) Peptide neuromodulation in invertebrate model systems. Neuron 76:82–97.

Thirumalai V, Marder E (2002) Colocalized neuropeptides activate a central pattern generator by acting on different circuit targets. J Neurosci 22:1874–1882.

Thirumalai V, Prinz AA, Johnson CD, Marder E (2006) Red pigment concentrating hormone strongly enhances the strength of the feedback to the pyloric rhythm oscillator but has little effect on pyloric rhythm period. J Neurophysiol 95:1762–1770.

Tohidi V, Nadim F (2009) Membrane resonance in bursting pacemaker neurons of an oscillatory network is correlated with network frequency. J Neurosci 29:6427–6435.

Tseng HA, Nadim F (2010) The membrane potential waveform of bursting pacemaker neurons is a predictor of their preferred frequency and the network cycle frequency. J Neurosci 30:10809–10819.

van den Pol AN (2012) Neuropeptide transmission in brain circuits. Neuron 76:98–115.

Wester JC, McBain CJ (2014) Behavioral state-dependent modulation of distinct interneuron subtypes and consequences for circuit function. Curr Opin Neurobiol 29:118–125.

White RS, Spencer RM, Nusbaum MP, Blitz DM (2017) State-dependent sensorimotor gating in a rhythmic motor system. J Neurophysiol 118:2806–2818.

Woods IG, Schoppik D, Shi VJ, Zimmerman S, Coleman HA, Greenwood J, Soucy ER, Schier AF (2014) Neuropeptidergic signaling partitions arousal behaviors in zebrafish. J Neurosci 34:3142–3160.

Xia XB, Mills SL (2004) Gap junctional regulatory mechanisms in the AII amacrine cell of the rabbit retina. Vis Neurosci 21:791–805.

Yamazoe-Umemoto A, Fujita K, Iino Y, Iwasaki Y, Kimura KD (2015) Modulation of different behavioral components by neuropeptide and dopamine signalings in non-associative odor learning of Caenorhabditis elegans. Neurosci Res 99:22–33.

Zhao S, Sheibanie AF, Oh M, Rabbah P, Nadim F (2011) Peptide neuromodulation of synaptic dynamics in an oscillatory network. J Neurosci 31:13991–14004.

